# Properties of Photosystem II lacking the PsbJ subunit

**DOI:** 10.1101/2021.09.04.458961

**Authors:** Alain Boussac, Julien Sellés, Marion Hamon, Miwa Sugiura

## Abstract

Photosystem II (PSII), the oxygen-evolving enzyme, consists of 17 trans-membrane and 3 extrinsic membrane proteins. Other subunits bind to PSII during assembly, like Psb27, Psb28, Tsl0063. The presence of Psb27 has been proposed (Zabret et al. 2021; Huang et al. 2021; Xiao et al. 2021) to prevent the binding of PsbJ, a single transmembrane α-helix close to the quinone Q_B_ binding site. Consequently, a PSII rid of Psb27, Psb28 and Tsl0034 prior to the binding of PsbJ would logically correspond to an assembly intermediate. The present work describes experiments aiming at further characterizing such a ΔPsbJ-PSII, purified from the thermophilic *Thermosynechococcus elongatus*, by means of MALDI-TOF spectroscopy, Thermoluminescence, EPR spectroscopy and UV-visible time-resolved spectroscopy. In the purified ΔPsbJ-PSII, an active Mn_4_CaO_5_ cluster is present in 60-70 % of the centers. In these centers, although the forward electron transfer seems not affected, the *Em* of the Q_B_/Q_B_^-^ couple increases by ≥ 120 mV thus disfavoring the electron coming back on Q_A_. The increase of the energy gap between Q_A_/Q_A_^-^ and Q_B_/Q_B_^-^ could contribute in a protection against the charge recombination between the donor side and Q_B_^-^, identified at the origin of photoinhibition under low light (Keren et al. 1997), and possibly during the slow photoactivation process.

## Introduction

Photosystem II (PSII), the water-splitting enzyme in cyanobacteria, algae and higher plants, is responsible for the production of the atmospheric O_2_ that is essential for aerobic organisms and it is the first step in the production of food, fibers and fossil fuels. The mature Photosystem II, in cyanobacteria, consists of 20 subunits (Umena et al. 2011; Suga et al. 2015) with 17 trans-membrane and 3 extrinsic membrane proteins (Roose et al. 2016). The PSII also binds 35 chlorophylls, 2 pheophytins, 2 hemes, 1 non-heme iron, 2 plastoquinones (Q_A_ and Q_B_), the Mn_4_CaO_5_ cluster, 2 Cl^-^, 12 carotenoids and 25 lipids (Umena et al. 2011; Suga et al. 2015). Most of the cofactors involved in the water oxidation and electron transfer bind to the reaction center subunits, PsbA (also called D1) and PsbD (also called D2).

The first step in the photosynthetic electron transfer chain is the absorption of a photon by the antenna chlorophyll. Then, the excitation is transferred to the photochemical trap that includes the four chlorophylls P_D1_, P_D2_, Chl_D1_ and Chl_D2_. After a few picoseconds following the light-absorption process a charge separation occurs forming the Chl_D1_^+^Phe_D1_^-^ and then the P_D1_^+^Phe_D1_^-^ radical pairs (Holzwarth et al. 2006; Romero et al. 2017). Then, P_D1_^+^ oxidizes Tyr_Z_, the Tyr161 of the D1 polypeptide, which in turn oxidizes the Mn_4_CaO_5_ cluster, see (Shen 2015, Cox et al. 2020) for reviews. The electron on Phe_D1_^-^ is then transferred to Q_A_, the primary quinone electron acceptor and then to Q_B_, the second quinone electron acceptor. Whereas Q_A_ is only singly reduced under normal conditions, Q_B_ accepts two electrons and two protons before to leave its binding site, reviewed in (de Causmaecker et al. 2019). The Mn_4_CaO_5_ cluster is sequentially oxidized following each charge separation so that it cycles through five redox states denoted S_*n*_, where *n* stands for the number of stored oxidizing equivalents (Kok et al. 1970; Joliot and Kok 1975). When the S_4_-state is formed, *i.e.* after the 3^rd^ flash of light given on dark-adapted PSII that is in the S_1_ state, the two water molecules bound to the cluster are oxidized, the O_2_ is released and the S_0_-state is reformed.

In cyanobacteria, in addition to the subunits involved in the cofactors binding (PsbA, PsbD, CP43, CP47, PsbE, PsbF), there are several small subunits consisting in only a transmembrane α-helix as PsbT, PsbM, PsbJ, PsbK, PsbL, PsbI, PsbX, PsbY, Psb30 (formerly Ycf12), *e.g.*(Kashino et al. 2002, 2007; Nowaczyk et al. 2012; Sugiura et al. 2012). Some roles have been attributed to these low molecular weight subunits either upon site directed mutagenesis, mainly for PsbT, or upon the deletion of the protein as for PsbM, PsbJ, Psb30, PsbK, PsbX, PsbZ.

It has been proposed that the interaction between Phe239 of PsbA and the PsbT subunit is required to restrict the movement of the DE loop of PsbA. In turn, the disruption of this interaction may perturb the binding of bicarbonate to the non-heme iron that could contribute to the signal for PSII to undergo a repair following photodamages (Forsman and Eaton-Rye 2021). Deletion of PsbM has been reported to mainly affect the Q_B_ environment (Uto et al. 2017). PSII depleted of Psb30 exhibited a lower efficiency under high light conditions and Psb30 favors the stabilization of the PSII complex (Sugiura et al. 2010a; Inoue-Kashino et al. 2011). Deletion of PsbK has been proposed to destabilize the association of PsbZ and Psb30 (Ycf12) with PSII complex and to alter the Q_B_ function (Iwai et al. 2010). PsbZ has been proposed to stabilize the binding of Psb30 (Takasaka et al. 2010). Deletion of PsbX has been shown to affect the PSII integrity in both *Arabidopsis thaliana* (Garcia-Cerdan et al. 2009) and *Synechocystis sp.* PCC 6803 (Funk 2000). In plant PSII, the PsbL subunit seems to prevent the back electron flow from the reduced plastoquinol pool thus protecting the PSII from photoinactivation (Ohad et al. 2004). In *Synechocystis sp.* PCC 6803, PsbL also influences forward electron transfer from Q_A_^-^ to Q_B_ (Luo et al. 2014).

The *psbJ* gene belongs to the *psbEFLJ* operon coding for the PsbE and PsbF subunits bearing the two histidine residues His23 and His24, respectively, which are the heme iron axial ligands of the Cyt*b*_559_ (Umena et al. 2011). The PsbJ subunit is close to PsbK and its possible involvement in the exchange of the plastoquinone has been discussed (Kaminskaya et al. 2007; Müh et al. 2012; van Eerden et al. 2017) with a role in the efficiency of forward electron flow following the charge separation process by affecting the Q_A_ and Q_B_ properties (Ohad et al. 2004; Regel et al. 2001). In *Synechocystis sp.* PCC 6803, double mutants lacking PsbJ and either PsbV or PsbO are unable to grow photoautotrophically (Choo et al. 2021).

In most of the deletion mutants, the observed phenotype is an alteration of the acceptor side. However, it is difficult to attribute these changes more to a specific role of the deleted subunits, rather than to perturbations in the overall structure of PSII. Cyanobacteria have several PsbA isoforms (Mulo et al. 2009; Sugiura and Boussac, 2014; Sheridan et al. 2020). In *Thermosynechococcus elongatus*, the deletion of the *psbJ* gene has different consequences with either PsbA1 or PsbA3 as the D1 protein. In PsbA3-PSII, the effects are minor whereas in the purified ΔPsbJ-PsbA1/PSII several other subunits including PsbY, PsbU, and PsbV are lacking (Sugiura et al. 2010b). In contrast, Psb27, Psb28 and Tsl0063, have been found to be associated to a proportion of the ΔPsbJ-PsbA1/PSII (Nowaczyk et al. 2012). These three proteins are known to be PSII assembly factors (Nowaczyk et al. 2006; Roose and Pakrasi 2008; Komenda et al. 2012; Liu et al. 2013; Huang et al. 2021; Zabret et al. 2021). It should be noted that Tsl0063 is named either Psb34 in (Zabret et al. 2021) or Psb36 in (Xiao et al. 2021). Structures of PSII corresponding to assembly intermediates have been recently solved by using cryo-EM by using different strategies (Zabret et al. 2021; Huang et al. 2021; Xiao et al. 2021). Upon deletion of the *psbJ* gene it became possible to isolate a “PSII-I” intermediate with Psb27, Psb28 and Tsl0063 bound and in which the major conformational change was a distortion of the Q_B_ binding site and the replacement of bicarbonate with glutamate as a ligand of the non-heme iron (Zabret et al. 2021). From a *psbV* deletion mutant a dimeric Psb27-PSII (Huang et al. 2021) and Psb28-PSII (Xiao et al. 201) have also been purified in which the dissociation of PsbJ occurs. Such changes are proposed to protect the PSII from damage during biogenesis until an active Mn_4_CaO_5_ cluster is assembled (Zabret et al. 2021; Xial et al. 2021).

Since the presence of Psb27 prevents the binding of PsbU and induces the dissociation of PsbJ and PsbY (Huang et al. 2021) and since PsbJ also seems to trigger the release of Psb28 (Zabret et al. 2021), the binding of PsbJ during the PSII assembly process very likely occurs *after* the release of Psb27 and Psb28. Consequently, a PSII without Psb27, Psb28, Tsl0063, and without PsbJ would logically also correspond to an assembly intermediate. In the present work, we describe the results of experiments aiming at further characterizing the ΔPsbJ-PsbA1/PSII purified from *Thermosynechococcus elongatus* by means of MALDI-TOF spectroscopy, Thermoluminescence, EPR spectroscopy and UV-visible time-resolved spectroscopy.

## Materials and Methods

### Samples used

The *Thermosynechococcus elongatus* strains used were; *i*) the *ΔpsbA_2_, ΔpsbA_3_* deletion mutant, referred to as either WT*1-PSII or PsbA1-PSII, *ii*) the *ΔpsbA_1_, ΔpsbA_2_* deletion mutant, referred to as either WT*3-PSII or PsbA3-PSII (Sugiura et al. 2010b), and *iii*) the ΔPsbJ-43H deletion mutant (Sugiura et al. 2010b) which has the three *psbA* genes but in which only the PsbA1-PSII is produced under the culture conditions used in this work, see thereafter. These strains were constructed from the *T. elongatus* 43-H strain that had a His_6_-tag on the carboxy terminus of CP43 (Sugiura and Inoue 1999). PSII purification was achieved as previously described (Sugiura et al. 2014). The final resuspending medium contained 1 M betaine, 15 mM CaCl_2_, 15 mM MgCl_2_, 40 mM Mes, pH 6.5 adjusted with NaOH.

### MALDI-TOF measurements

All reagents and solvents were purchased from Sigma-Aldrich (Saint Quentin-Fallavier, France) with the highest purity available. Peptides and protein used for calibration were purchased from LaserBio Labs (TOF Mix) and Sigma (equine apomyoglobin), respectively. For intact mass analysis, 1 µL of PSII complex prepared at a concentration of ∼100 µg Chl /mL in the medium mentioned above was mixed with 2 µL of a saturated solution of sinapinic acid in 60/0.1 acetonitrile/trifluoroacetic acid. Two microliters of this premix were spotted onto the sample plate and allowed to dry under a gentle air stream. Spectra were acquired in positive reflectron and linear mode on an Axima Performance MALDI-TOF/TOF mass spectrometer (Shimadzu, Manchester, UK) with a pulse extraction fixed at 4000 for 3000-10000 m/z range and at 10000 for 6000-20000 m/z range acquisitions. All spectra were externally calibrated using a homemade calibrant mixture prepared by mixing 1 µL of 50 µM apomyoglobine in water with 2 µL of TOF Mix solution containing ACTH [7-38] peptide at a concentration of 6 µM.

### UV-visible time-resolved absorption change spectroscopy

Absorption changes measurements have been performed with a lab-built spectrophotometer (Béal et al. 1999) in which the absorption changes were sampled at discrete times by short analytical flashes. These analytical flashes were provided by an optical parametric oscillator (Horizon OPO, Amplitude Technologies) pumped by a frequency tripled Nd:YAG laser (Surelite II, Amplitude Technologies), producing monochromatic flashes (355 nm, 2 nm full-width at half-maximum) with a duration of 5 ns. Actinic flashes were provided by a second Nd:YAG laser (Surelite II, Amplitude Technologies) at 532 nm, which pumped an optical parametric oscillator (Surelite OPO plus) producing monochromatic saturating flashes at 695 nm with the same pulse-length. The two lasers were working at a frequency of 10 Hz. The time delay between the laser delivering the actinic flashes and the laser delivering the detector flashes was controlled by a digital delay/pulse generator (DG645, jitter of 1 ps, Stanford Research). The path-length of the cuvette was 2.5 mm.

The samples were diluted in 1 M betaine, 15 mM CaCl_2_, 15 mM MgCl_2_, and 40 mM Mes (pH 6.5). PSII samples were dark-adapted for ∼ 1 h at room temperature (20–22°C) before, when indicated, the addition of 0.1 mM phenyl *p*–benzoquinone (PPBQ) dissolved in dimethyl sulfoxide. The chlorophyll concentration of all the samples was ∼ 25 µg of Chl/mL. After the ΔI/I measurements, the absorption of each diluted batch of samples was precisely controlled to avoid errors due to the dilution of concentrated samples and the ΔI/I values were normalized to *A*_673_ = 1.75, with ε ∼ 70 mM^-1^·cm^-1^ at 674 nm for dimeric PSII (Müh and Zouni 2005).

### EPR spectroscopy

X-band cw-EPR spectra were recorded with a Bruker Elexsys 500 spectrometer equipped with a standard ER 4102 (Bruker) X-band resonator, a Bruker teslameter, an Oxford Instruments cryostat (ESR 900) and an Oxford ITC504 temperature controller. Flash illumination at room temperature was provided by a neodymium:yttrium–aluminum garnet (Nd:YAG) laser (532 nm, 550 mJ, 8 ns Spectra Physics GCR-230-10). Illumination at 198 K with visible light was done in a non-silvered Dewar filled with ethanol cooled with dry ice for approximately 5-10 seconds with a 800 W tungsten lamp filtered by water and infrared cut-off filters. Illumination at 4.2 K was done in the EPR cavity using a low-voltage halogen lamp (24V, 250W, Philips Type 13163) filtered by water and infrared cut-off filters. After a 1 h dark-adaptation at room temperature the samples were frozen in the dark to 198 K in a dry ice ethanol bath and then transferred to 77 K in liquid N_2_. Prior to the recording of the spectra the samples were degassed at 198 K.

### Thermoluminescence measurements

Thermoluminescence (TL) curves were measured with a lab–built apparatus (Ducruet 2003; Ducruet and Vass 2009). PSII samples were diluted in 1 M betaine, 40 mM MES, 15 mM MgCl_2_, 15 mM CaCl_2_, pH 6.5 and then dark-adapted for at least 1 h at room temperature. When used, DCMU (100 µM, final concentration), dissolved in ethanol, was added after the dark-adaptation. Flash illumination was given at −10°C by using a saturating xenon flash. Two reasons justifies the choice of this temperature. Firstly, with the resuspending medium used, the freezing/melting of the samples occurs at ∼ −15°C. Since the frozen samples are strongly diffusing, the flash illumination was done at a temperature slightly above −15°C. Secondly, an artefact occurs in the melting region which makes difficult the detection of small signals. It has been checked after the dilutions that the PSII samples had the same OD at 673 nm equal to 0.70 *i.e.* ∼ 10 µg Chl/mL.

The measurements have been done on two different ΔPsbJ-PSII preparations with at least two measurements on each PSII samples except for the period 4 oscillation experiment at 291 nm done on only one PSII preparation. In addition to MALDI-TOF analysis, the protein content of the ΔPsbJ-PSII has been previously monitored with SDS-page and the oligomeric state of PSII was also analyzed by gel permeation (Sugiura et al. 2010b). Most of the ΔPsbJ-PSII, although not all, was in monomeric form and there was no evidence for the presence of any of the 3 proteins Psb28, Psb27 and Tsl0063 in significant amounts. For the spectroscopic measurements all samples were dark-adapted for one hour at room temperature before the measurements for allowing a full decay of S_2_ and S_3_ into S_1_ (Sugiura et al. 2004).

## Results

Before to study the effects of the PsbJ deletion some control experiments were performed. Indeed, it has been reported (Nowaczyk et al. 2012) that in *T. elongatus* cells having all the *psbA1, psbA2* and *psbA3* genes, and the *psbJ* gene inactivated by insertion of a cassette, PsbA1 was replaced by PsbA3 in the ΔPsbJ-PSII. Although this substitution was not observed in the most recent work done by the same group (Zabret et al. 2021) it seemed important to us to verify that the D1 subunit was indeed PsbA1 in our ΔPsbJ-43H/PSII which also contains the 3 genes, *psbA1, psbA2* and *psbA3*. For that, we have recorded the electrochromic blue shift undergone by Phe_D1_ in the Q_X_ spectral region upon the formation of Q_A_^-^. This electrochromic C-550 band shift is well known to be red shifted by ∼ 3.0 nm from 544 nm in PsbA1-PSII to 547 nm in PsbA3-PSII. This shift reflects a hydrogen bond from the 13^1^-keto C=O group of Phe_D1_ stronger with the carboxylate group of E130 in PsbA3-PSII than with the amine group of Q130 in PsbA1-PSII (Merry et al. 1998; Cuni et al. 2004; Hughes et al. 2010; Shibuya et al. 2010).

Figure 1 shows the C–550 bandshift with PsbA1-PSII (black spectrum), in PsbA3-PSII (blue spectrum) and in ΔPsbJ-43H/PSII (red spectrum). For these measurements, the samples were first dark–adapted for one hour at room temperature. Then, the absorption changes were measured 15 µs after each actinic flash in a series in the presence of PPBQ. The data points are the average of the individual absorption changes induced by the 2^nd^ to 7^th^ flashes. In the ΔPsbJ-43H/PSII the electrochromic band shift was similar to that in the PsbA1/PSII sample, showing unambiguously that the D1 protein is PsbA1 in the ΔPsbJ-43H/PSII.

**Figure 1:**
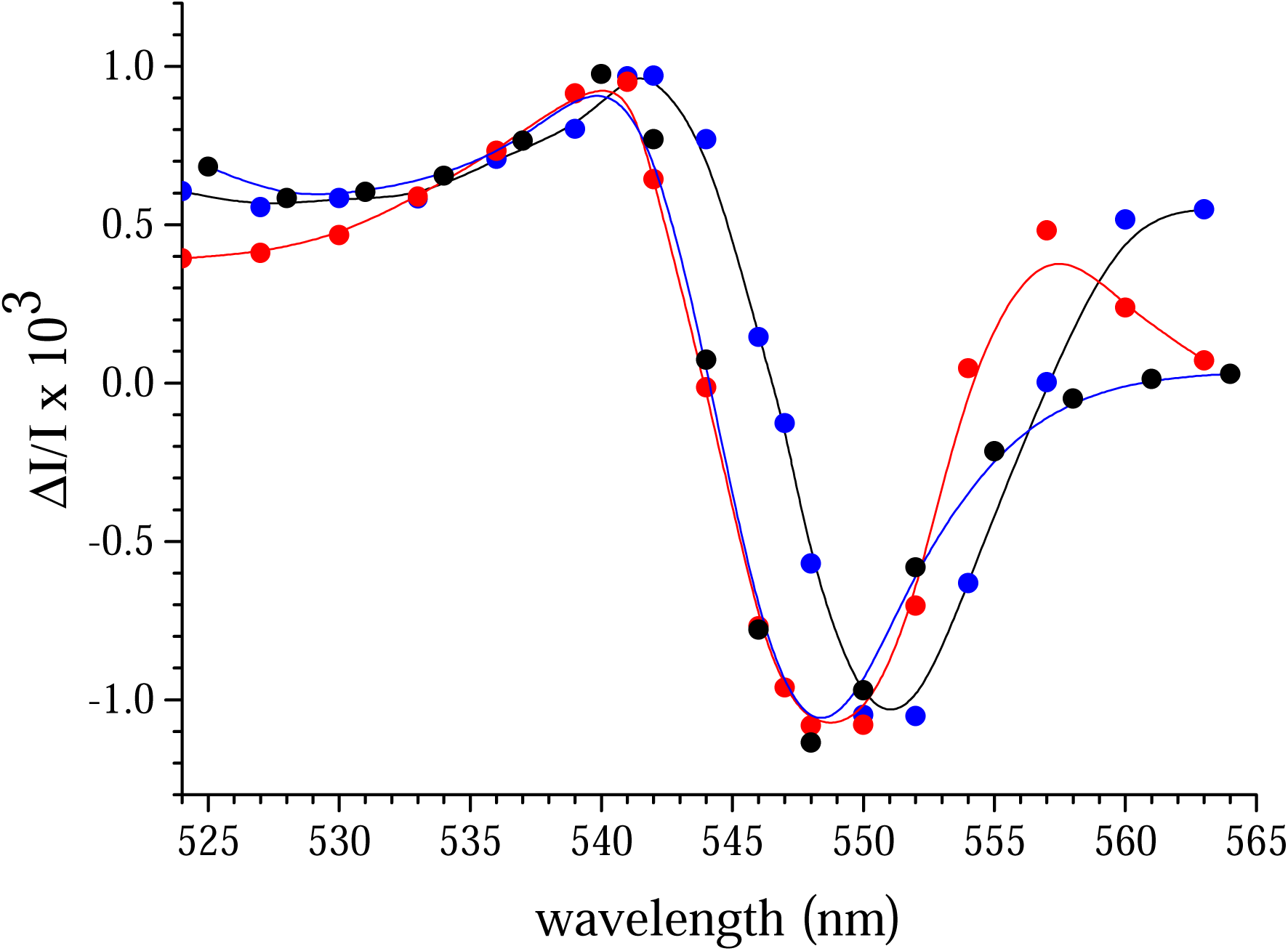
Light-induced difference spectra around 545 nm. The flash-induced absorption changes were measured in PsbA1/PSII (black spectrum), ΔPsbJ-43H/PSII (red spectrum) and PsbA3-PSII (blue spectrum). The data points are the average of the ΔI/I values detected 15 μs after the 2^nd^ to 7^th^ actinic flashes given to dark-adapted PSII. After dark adaptation for 1 h at room temperature, 100 μM PPBQ (dissolved in dimethyl sulfoxide) was added to the samples. The amplitude of the spectra were normalized to a Chl concentration corresponding to OD_673nm_=1.75.

Secondly, the polypeptide content was analysed in WT*1-PSII and ΔPsbJ-43H/PSII with MALDI-TOF spectroscopy. Panel A in Figure 2 shows the *m/z* region from 3500 to 6500. The inset in Panel A shows the *m/z* region corresponding to PsbM (4009 Da calc.) and PsbJ (4002 Da calc.) by a using the reflection mode. The different peaks for one protein correspond to the different proportions of ^13^C in the proteins starting from no ^13^C (100% ^12^C) to 1, 2, 3, and so on, ^13^C per protein. The two peaks for PsbJ spaced by ∼16 *m/z* likely correspond to an oxygen adduct for the larger *m/z* value, also seen in earlier reports as in (Sugiura et al. 2010b;Nowaczyk et al. 2012). Panel B in Figure 2 shows the *m/z* region from 6500 to 16000. The PsbJ subunit, as expected, is missing in the ΔPsbJ-43H/PSII (Sugiura et al. 2010b; Nowaczyk et al. 2012). In ΔPsbJ-43H/PSII, other subunits are missing as PsbY, PsbU, PsbV (the Cyt*c*_550_) as previously observed. However, PsbM and PsbF are now detected in 43H/ΔPsbJ-PSII exhibiting a peak with an amplitude comparable to that in WT*1-PSII and with a single peak for PsbM while two peaks (native and formylated) were previously observed in (Sugiura et al. 2010b). The reasons for such a difference between the two observations is at present unclear. A reexamination of the raw data in (Sugiura et al. 2010b) indicates that at that time we did not pay attention to the possible presence of Psb28 in a very small proportion of our ΔPsbJ-43H/PSII. Indeed, a very small peak at *m/z* ∼ 12787 was present (Sugiura et al. 2010b). Panel B in Figure 2 also shows a very small peak at 12787 that could well correspond to Psb28. However, the amplitude of this peak is close to the limit of the detection. A small peak detected at *m/z*=13424 in some traces could correspond to Psb27 (not shown). A peak at *m/z*=5936, not attributed in (Sugiura et al. 2010b) could originate from Tsl0063 recently detected in ΔPsbJ-PsbA1/PSII (Zabret et al. 2021). This protein is however not detected in Figure 2. The MALDI-TOF data described above in comparison with those in literature show that upon the deletion of PsbJ in PsbA1-PSII, the PSII composition may slightly vary from prep to prep. If present, the 3 proteins Psb27, Psb28 and Tsl0063 are in a so minor proportion of the ΔPsbJ-43H/PSII studied here that this proportion will not contribute significantly to the results described thereafter. We cannot eliminate the possibility that these 3 proteins are removed during the PSII purification. This, however, does not modify the fact that the studied ΔPsbJ-43H/PSII does not bind Psb27, Psb28 and Tsl0063.

**Figure 2:**
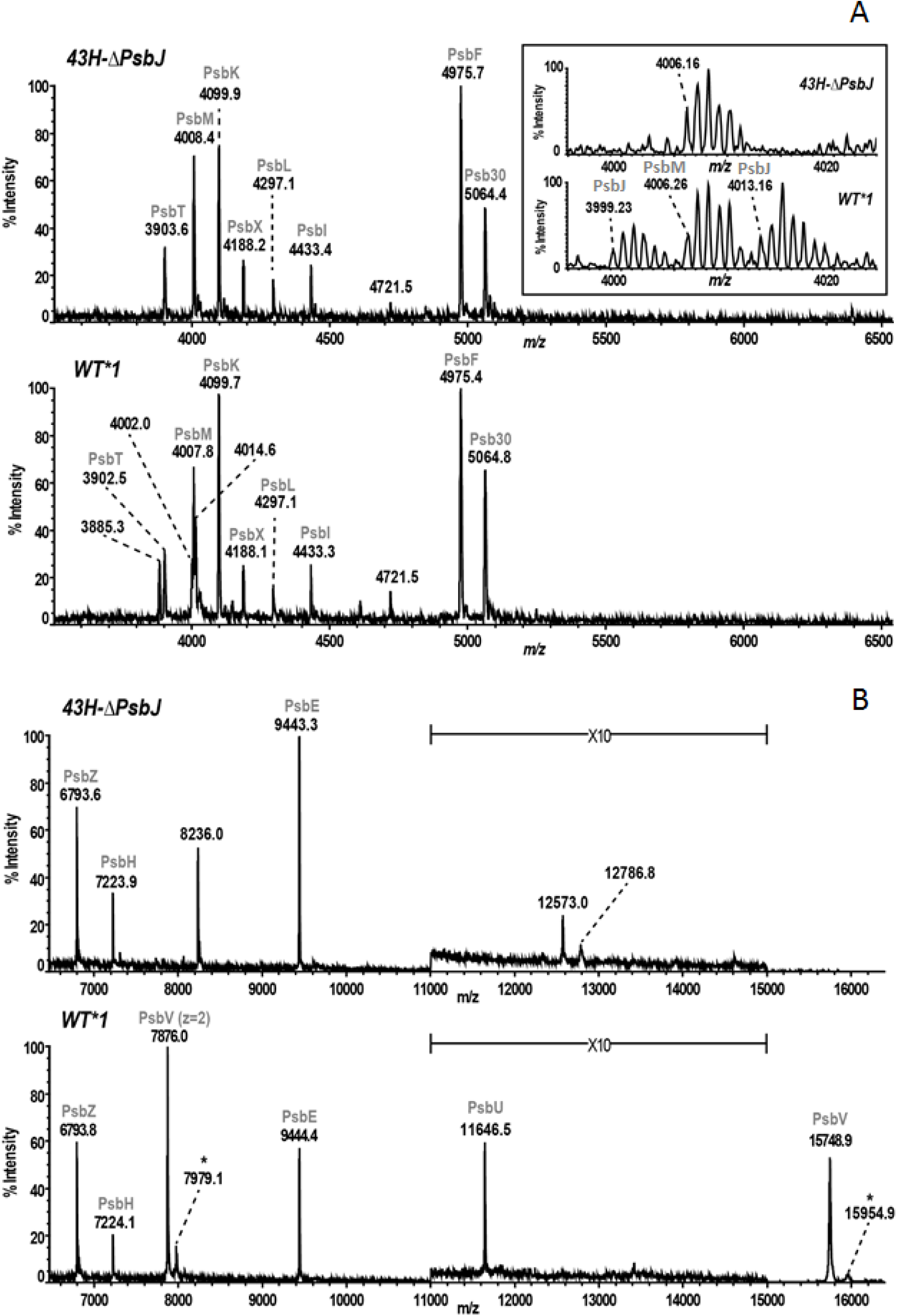
MALDI-TOF MS profiling of PSII subunits. (A) Linear mode MALDI-TOF spectra of subunits from ΔPsbJ-43H/PSII (upper panel) and WT*1-PSII (lower panel) strains. For presentation, both spectra were internally recalibrated on known peaks of PSII subunits using their theoretical average masses (formylated PsbT, *m/z* = 3902.67 Da; PsbK, *m/z* = 4099.88 Da; PsbL, *m/z* = 4297.02 Da; acetylated PsbF, *m/z* = 4975.66 Da; formylated PsbZ, *m/z* = 6792.18 Da; PsbE, *m/z* = 9441.53Da) according to (Sugiura et al. 2010a 2010b, Boussac et al. 2013, Nowaczyk et al. 2012). In the inset, zoom of the 3090-4025 *m/z* range of high-resolution MALDI-TOF spectra from ΔPsbJ-43H/PSII mutant (upper panel) and WT*1-PSII wild-type (lower panel) strains acquired in reflection mode. Both spectra were internally recalibrated on known mono-isotopic peaks of PSII subunits using their theoretical mono-isotopic masses (formylated PsbT, *m/z* = 3900.09 Da; PsbK, *m/z* = 4097.32 Da; PsbL, *m/z* = 4294.32 Da; acetylated PsbF, *m/z* = 4972.61 Da. (B) Linear mode MALDI-TOF spectra of ΔPsbJ-43H/PSII mutant (upper panel) and WT*1/PSII (lower panel) strains. In the upper panel which shows the 11000-15000 *m/z* range for the ΔPsbJ-43H/PSII mutant spectrum, the amplitude was magnified 10 times. (*) this 206 Da mass shift could correspond to farnesyl adduct find in mono-charged and discharged PsbV. There is no peak at 16472 *m/z* which suggests that the peak at 8236 *m/z* does not correspond to a double charged ion. Instead, it could correspond to a contamination by the subunit c of the ATP synthase (Suhai et al. 2008).

In *Synechocystis* 6803, the deletion of PsbJ has been proposed to alter both the forward electron flow from Q_A_^-^ to the plastoquinone pool and the back reaction between Q_A_^-^ and the oxidized Mn_4_CaO_5_ cluster (Regel et al. 2001). These conclusions were done in part from thermoluminescence experiments (Rutherford et al. 1982) in whole cells. Here, similar measurements have been done but in purified WT*1-PSII and ΔPsbJ-43H/PSII. Figure 3 shows the TL curves after 1 flash given at −10°C in WT*1-PSII, black traces, and ΔPsbJ-43H/PSII, red traces. In Panel A, the TL measurements were done without any addition whereas in Panel B they were done in the presence of DCMU. The TL curves in WT*1-PSII arising from the S_2_Q_B_^−^ charge recombination in Panel A with a peak temperature at ∼43°C and from the S_2_Q_A_^−^/DCMU charge recombination in Panel B with a peak temperature at ∼13°C are those expected in *T. elongatus* with an heating rate of 0.4°C/s. In ΔPsbJ-43H/PSII, no signal was detected both in Panels A and B. The very small signal around 0°C in Panels A and B is very similar in the absence and the presence of DCMU and therefore cannot arise from the S_2_Q_B_^−^ and S_2_Q_A_^−^/DCMU charges recombinations studied here. The TL results in Figure 3 differ significantly from those in *Synechocystis* whole cells where in the ΔPsbJ mutant a small TL signal was detected at a slightly lower temperature for the S_2_Q_B_^−^ charge recombination and at a slightly higher temperature for the S_2_Q_A_^−^/DCMU charge recombination (Regel et al. 2001). However, the amplitude of a TL signal depends on the proportion of centers with Q_B_^-^ in the dark-adapted state, something that is difficult to control in whole cells. The lack of a TL signal in ΔPsbJ-43H/PSII both with and without DCMU may be explained if 100% of the centers are in the Q_B_^-^ state upon the dark adaptation. As this will be shown later, that is not the case in the ΔPsbJ-43H/PSII studied here and we have to find another explanation.

**Figure 3:**
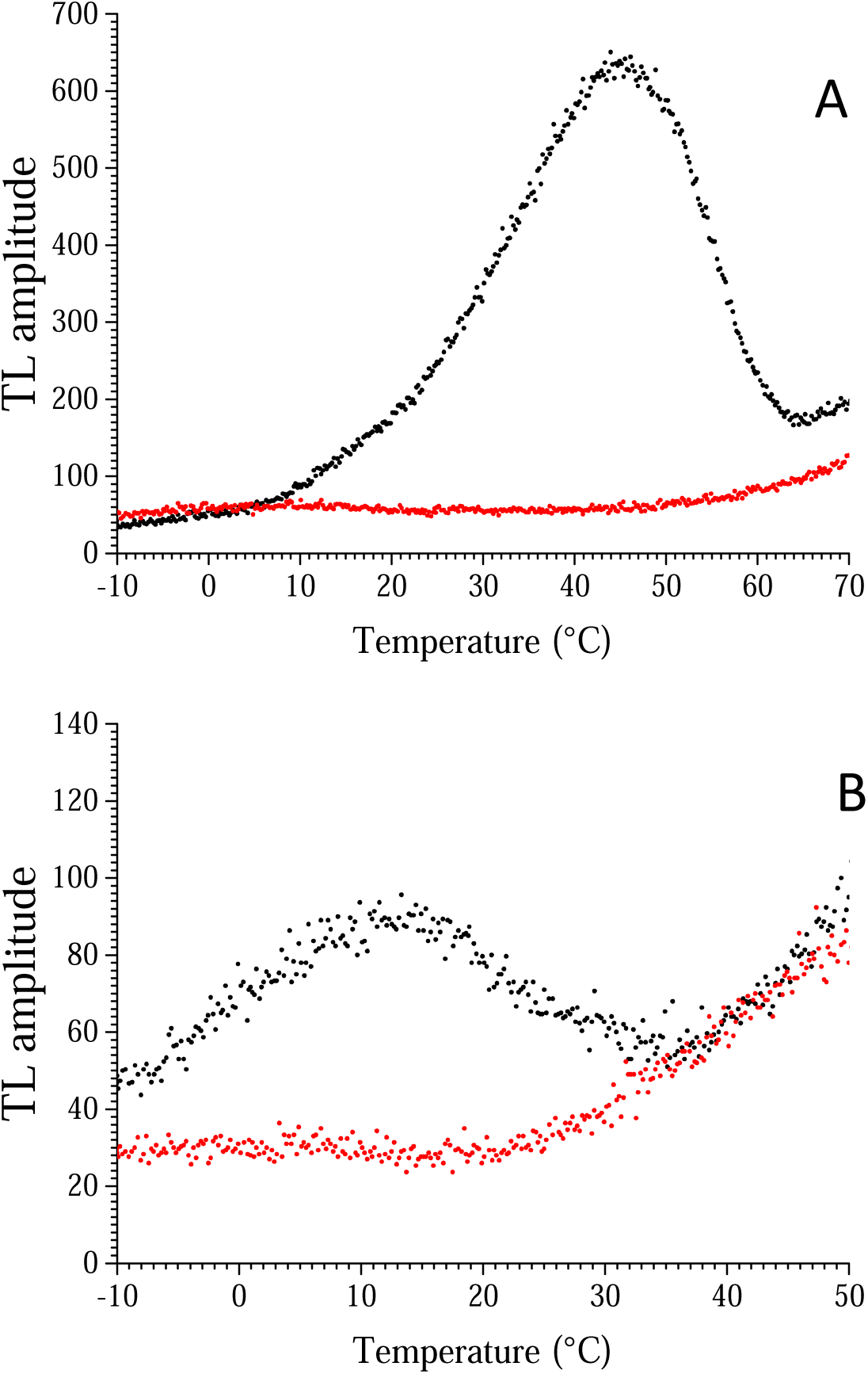
Thermoluminescence curves after one flash given at −10°C either without any addition (Panel A) or in the presence of DCMU (Panel B). Black curve, WT*1/PSII; red curve, ΔPsbJ-43H/PSII. The concentration of the samples was adjusted exactly to OD_673nm_=0.7 (∼10 µg Chl/mL) before to be dark-adapted at room temperature for at least 1 h. In Panel B, the final concentration of DCMU, dissolved in ethanol, was 100 µM. After the addition of DCMU, the samples were immediately loaded into the cuvette in total darkness. The heating rate was 0.4°C/s.

The first simple explanation is that the radiative charge recombination in the ΔPsbJ-43H/PSII occurs either *i*) at a temperature much above a bearable temperature for PSII (*i.e.* >> 60°C) or *ii*) at a temperature much lower than 0°C. In the first case, the *Em* of the Q_A_/Q_A_^-^couple, upon the deletion of PsbJ, would be strongly increased and, in the second case, strongly decreased. The second possibility could be a situation in which the Mn_4_CaO_5_ cluster in the ΔPsbJ-43H/PSII is inactive or lost because we know that under continuous illumination the activity is significantly decreased (Sugiura et al. 2010b).

Before to go further in the interpretation of the TL experiments we have therefore performed some EPR measurements for clarifying the situation.

Figure 4 shows the EPR spectra recorded at 8.6 K in dark-adapted samples (black spectra) and after a continuous illumination at 198 K (red spectra). The blue spectra are the light-*minus*-dark spectra. At 198 K, the electron transfers from Q_A_^-^ to Q_B_ and from Q_A_^-^ to Q_B_^-^ are blocked so that only one charge separation may occur (Fufezan et al. 2005). In WT*1-PSII, the black spectrum is very similar to the spectrum recorded in PsbA3-PSII under the same conditions, see (Boussac et al. 2011) for a detailed description of the spectra. The signal between 3600 and 5000 gauss originates from Fe^2+^Q_B_^-^ (Fufezan et al. 2005; Boussac et al. 2011; Sedoud et al. 2011). After illumination at 198 K for ∼ 5-10 s, the S_2_ multiline signal was formed together with a signal at *g* = 1.6 (∼ 4100 gauss). The *g* = 1.6 signal originates from the Q_A_^-^Fe^2+^Q_B_^-^ state. The Q_A_^-^Fe^2+^ signal formed in centers with Q_B_ oxidized prior to the illumination is difficult to detect in the presence of the multiline signal. A careful examination of the blue spectrum in the *g* = 6 to *g* = 8 region indicates that the non-heme iron was oxidized in a very small fraction of the dark-adapted centers and was reduced upon the 198 K illumination (Boussac et al. 2011). The spectrum in the dark-adapted ΔPsbJ-43H/PSII also exhibits a Fe^2+^Q_B_^-^ signal with a somewhat smaller amplitude (see also Figure 7) than in WT*1-PSII. This observation rules out the hypothesis that the lack of a TL signal is due to centers with 100% Q_B_^-^ in the dark-adapted state. In agreement with the MALDI-TOF data above, and as previously reported (Sugiura et al. 2010b), the EPR also shows that the Cyt*c*_550_ (PsbV) is missing in the purified PSII as evidenced by the much smaller signals at *g* ∼ 3 and *g* ∼ 2.2 (3000 gauss) corresponding to the *g*_z_ and *g*_y_ resonances of cytochrome signals, respectively (the *g*_x_ feature is difficult to detect under the conditions used for the recording of these spectra). We have nevertheless previously shown that PsbV was detectable in the ΔPsbJ-43H/thylakoids (Sugiura et al. 2010b). Cyt*b*_559_ is detected because it is in the low potential oxidized form in ΔPsbJ-43H/PSII (Sugiura et al. 2010). The detection of a signal at *g* = 6 (∼1100 gauss) indicates that a proportion of Cyt*b*_559_ is also present in an oxidized high spin (*S* = 5/2) configuration. Importantly, the S_2_ multiline signal is detectable upon the 198 K illumination with an amplitude close to 60-70 % of that in WT*1-PSII. Therefore, the total absence of a TL signal in this sample is also not due to an inactive or missing Mn_4_CaO_5_ cluster. The proportion of ΔPsbJ-43H/PSII in which the Mn_4_CaO_5_ is able to progress to the S_2_ state upon an illumination at 198 K is higher than the O_2_ evolution activity found in this PSII that is about 30% (Sugiura et al. 2010b). However, it is well known that the O_2_ activity under continuous illumination does not necessarily correlate with the proportion of active centers. For example, upon the substitution of Ca^2+^ for Sr^2+^ the O_2_ activity under continuous illumination is decreased by a least of factor of 2 whereas all the centers are fully active (Ishida et al. 2008). In an additional control experiment (not shown) aiming at following the period four oscillation measured at 291 nm and 100 ms after each flash of a series (Lavergne 1991; Ishida et al. 2008) it has been found that in the ΔPsbJ-43H/PSII the miss parameter is close to 20% instead of ∼ 8% in the PsbA1-PSII.

**Figure 4:**
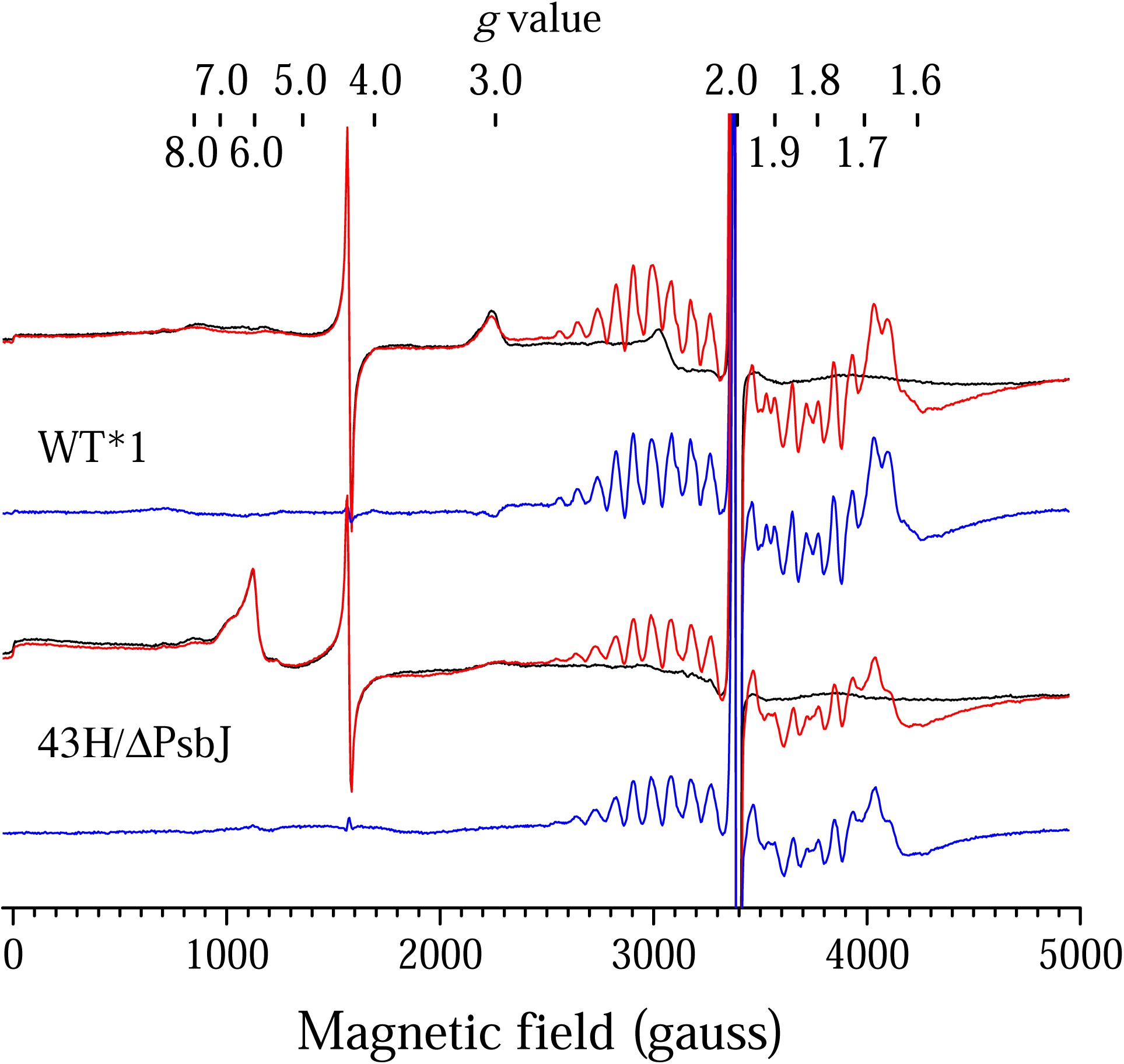
EPR spectra recorded in WT*1-PSII and ΔPsbJ-43H/PSII. The spectra were recorded after dark-adaptation for 1 hour at room temperature (black spectra) and after a continuous illumination at 198 K (red spectra). The blue spectra are the light-*minus*-dark spectra. The concentration was 1.1 mg Chl/mL. Instrument settings: temperature, 8.6 K; modulation amplitude, 25 G; microwave power, 20 mW; microwave frequency, 9.4 GHz; modulation frequency, 100 kHz.

The remaining possible explanations for the lack of a TL signal in ΔPsbJ-43H/PSII are either *i*) an inefficient charge separation after one flash illumination in contrast to the continuous illumination at 198 K, and/or *ii*) a too fast, even at −10°C, or too slow charge recombination in the temperature range probed by the TL thus preventing the detection of the signal. These possibilities have been tested in the EPR experiments reported in Figure 5 and Figure 6.

**Figure 5:**
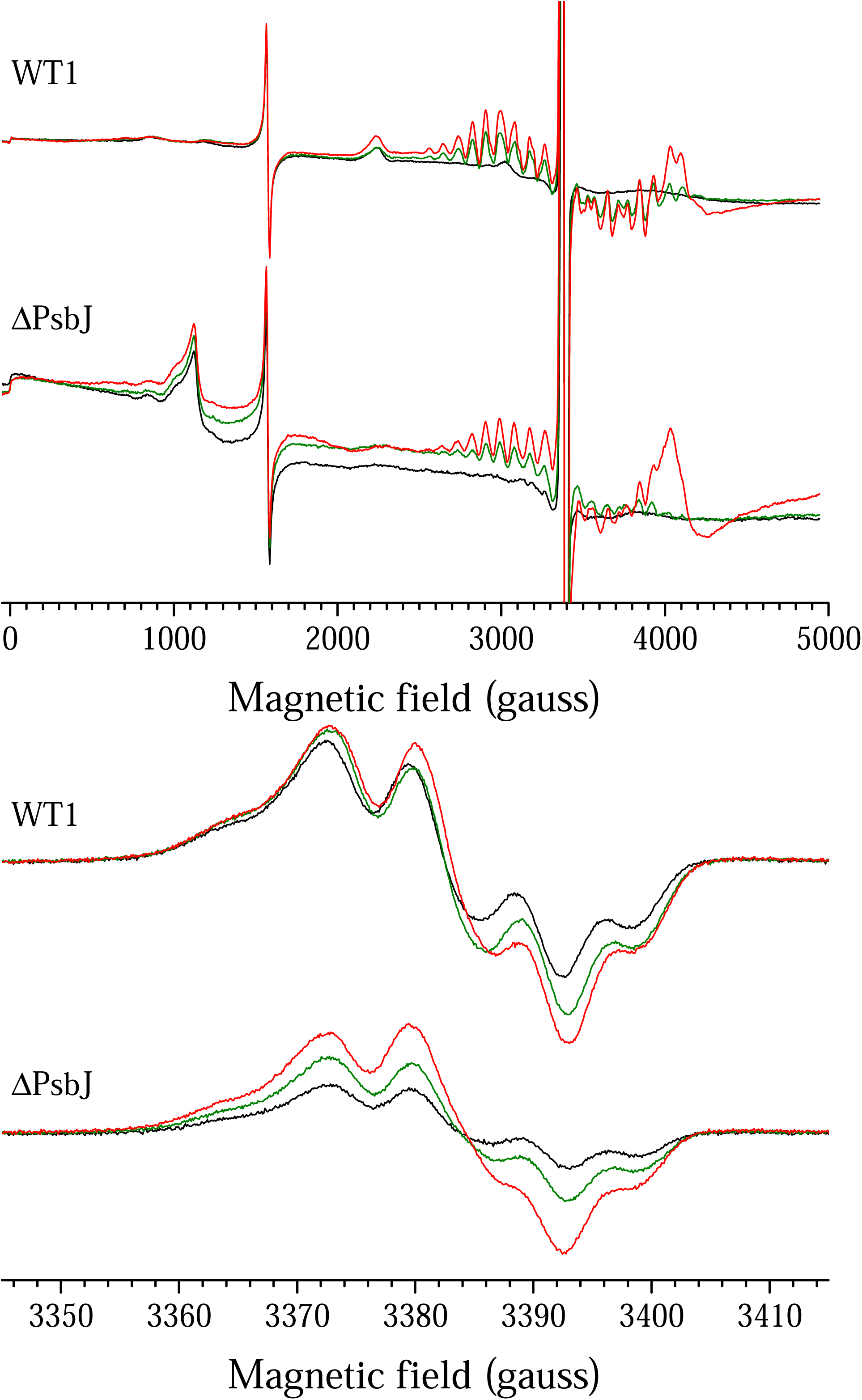
EPR spectra recorded in WT*1/PSII and ΔPsbJ-43H/PSII. The spectra were recorded after dark-adaptation for 1 hour at room temperature (black spectra), after illumination by one flash at room temperature (green spectra) and after a further continuous illumination at 198 K (red spectra). The concentration was 1.1 mg Chl/mL. Instrument settings: temperature, 8.6 K; microwave frequency, 9.4 GHz; modulation frequency, 100 kHz. Modulation amplitude, 25 G and microwave power, 20 mW in Panel A and modulation amplitude, 2.8 G and microwave power, 2 µW in Panel B. In the conditions used for the recording of the Tyr_D_ ^•^ spectra, the microwave power is still slightly saturating so that an increase in the relaxation properties upon the formation of S_2_ induces an increase of the signal amplitude (Styring and Rutherford 1988). This effect is larger in the negative part of the signal and is less when using a smaller modulation frequency (not shown) which is indicative of a rapid-passage effect (Styring and Rutherford 1988).

**Figure 6:**
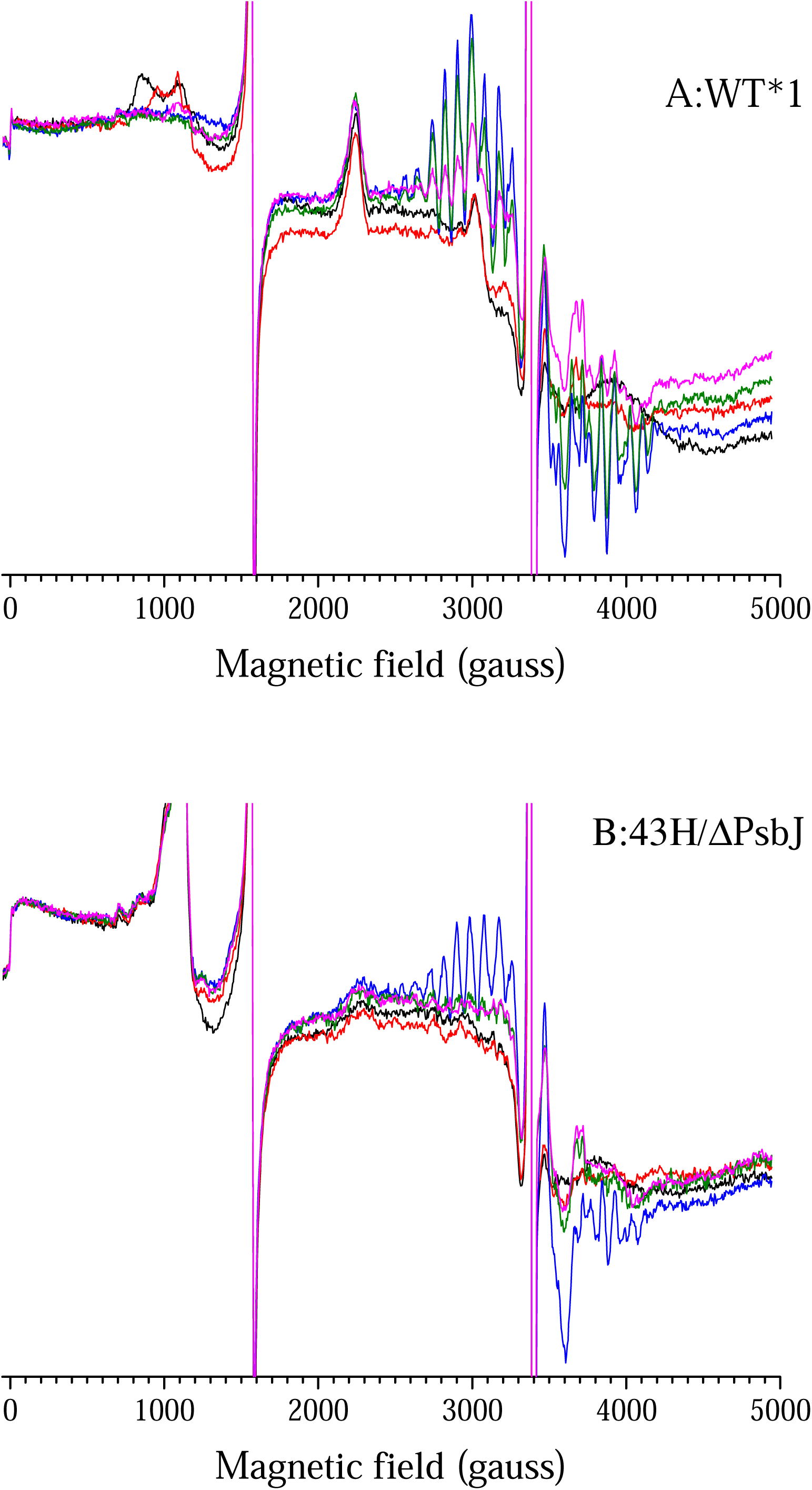
EPR spectra recorded in WT*1-PSII (Panel A) and ΔPsbJ-43H/PSII (Panel B). The spectra were recorded after dark-adaptation for 1 hour at room temperature (black spectra) and after the addition (100 µM) dissolved in ethanol (red spectra). Then, the blue spectra were recorded after an illumination at 198 K. The green spectra were recorded after a brief (1-2 s) warming of the samples at 0°C and the spectra in magenta were recorded after a second brief warming at 20°C. The concentration was 1.1 mg Chl/mL. Instrument settings: temperature, 8.6 K; microwave frequency, 9.4 GHz; modulation frequency, 100 kHz; modulation amplitude, 25 G; microwave power, 20 mW.

Panel A in Figure 5 shows the magnetic field range in which most of the EPR signals are detectable except Tyr_D_^•^. The Tyr_D_^•^ signal is shown in Panel B. The black spectra were recorded in dark-adapted PSII and they are, of course, similar to those described in Figure 4. The green spectra have been recorded after one saturating flash given at room temperature (∼20-22 °C) followed by an as-fast-as possible freezing of the sample (∼2 s) in a dry ice bath at 198 K. Then, the red spectra were recorded after a further continuous illumination at 198 K following the one flash illumination. In WT*1-PSII, the difference in the amplitude of the S_2_ multiline between the green and red spectra is mainly due to the proportion of centers in which the S_0_Tyr_D_^•^ to S_1_Tyr_D_^•^ transition occurs with the one flash illumination. As the Tyr_D_^•^ signal (Panel B) does not vary significantly, this suggests that the proportion of centers in which the S_1_Tyr_D_ to S_1_Tyr_D_^•^ transition occurs is negligible (Styring and Rutherford 1987).

In contrast, in ΔPsbJ-43H/PSII, upon the illumination at 198 K, the increase of the S_2_ multiline and Tyr_D_^•^ signals was much more pronounced than in WT*1-PSII. This results shows that in dark-adapted ΔPsbJ-43H/PSII there is a proportion of centers in the S_1_Tyr_D_ state and a larger proportion of centers in the S_0_Tyr_D_^•^ state than in WT*1-PSII. Since the proportion of centers in the Q_A_Fe^2+^Q_B_^-^ state is smaller in the ΔPsbJ-43H/PSII than in WT*1-PSII after the dark adaptation, this ratio is inversed after the one flash illumination. Consequently, the illumination at 198 K is expected to induce a larger Q_A_^-^Fe^2+^Q_B_^-^ in ΔPsbJ-43H/PSII than in WT*1-PSII and that is indeed what is observed here. It should be mentioned that, normally, an illumination at 198 K is unable to oxidize Tyr_D_ in active centers. The increase seen in Panel B therefore likely occurs in the proportion of inactive ΔPsbJ-43H/PSII. The important result in this experiment is that the one-flash illumination is able to induce the S_2_ state in the ΔPsbJ-43H/PSII although to a slightly less extent than in WT*1-PSII due to a higher proportion of centers in the S_1_Tyr_D_ and S_0_Tyr_D_^•^ states and also to a higher miss parameter. The stability of the S_2_ state has then been monitored in an experiment corresponding to the TL experiment in the presence of DCMU.

In Figure 6, the black spectra were recorded in dark-adapted PSII without any addition and the red spectra after the addition of DCMU. Two observations can be made here. Firstly, in both WT*1-PSII and ΔPsbJ-43H/PSII, the addition of DCMU resulted in the formation of Q_A_^-^Fe^2+^/DCMU to the detriment of Fe^2+^Q_B_^-^ (Velthuys 1981). In both samples, Q_A_^-^ Fe^2+^/DCMU exhibited either the *g* = 1.9 signal or the *g* = 1.82 signal (Rutherford et al. 1983, Rutherford and Zimmermann 1984), see below for a better description of the signals in Figure 6. Secondly, in WT*1-PSII, the addition of DCMU also modified the non-heme iron signal as previously observed (Diner and Petrouleas 1987). The blue spectra were then recorded after a continuous illumination at 198 K. The S_2_ multiline signal was formed in the open centers, *i.e.* in those with no Q_B_^-^ prior to the addition of DCMU. The larger Q_A_^-^Fe^2+^ signal at *g* = 1.9 in ΔPsbJ-43H/PSII than in WT*1-PSII have two possible explanations: *i*) a lower amount of Q_B_^-^ before the addition of DCMU and *ii*) a smaller proportion of oxidized non-heme iron. Indeed, the oxidation of Q_A_^-^ by the oxidized non-heme iron results in the disappearance of the non-heme iron signal is WT*1-PSII. Unfortunately, in ΔPsbJ-43H/PSII, the presence of the high spin Cyt*b*_559_ signal does not allow the detection of the non-heme iron signal.

After the illumination at 198 K, the samples were immersed for 1 to 2 seconds, in total darkness, in an ethanol bath at 0°C and immediately refrozen in a dry ice bath at 198 K and the green spectra were recorded. In WT*1-PSII, the brief passage at 0°C induced almost no change in the amplitude of the S_2_ multiline signal. In contrast, in ΔPsbJ-43H/PSII, almost all the S_2_ multiline signal and the *g* = 1.9 quinone signal disappeared thus showing that the recombination in the S_2_Q_A_^-^/DCMU state is very fast at 0°C in this sample. This very likely explains the lack of a TL signal in Figure 3B. The proportion of the *g* = 1.9 signal which decayed during the warming at 0°C appears larger than the proportion the *g* = 1.82 signal which decayed. This suggests that the recombination was more efficient with the quinone in the *g* = 1.9 state than in the *g* = 1.82 state (Demeter al. 1993; Rutherford, personal communication). Finally, the samples were immersed, in the dark, in an ethanol bath at 20°C and immediately refrozen in a dry ice bath at 198 K and the spectra in magenta were recorded. In WT*1-PSII, the S_2_ multiline now decreased significantly as expected from the peak temperature observed in this sample. In ΔPsbJ-43H/PSII, the spectrum was not significantly different from that one after the thawing at 0°C. The recording of the Tyr_D_^•^ spectra as in Panel B of Figure 5 showed again that Tyr_D_^•^ could be induced at 198 K and that a decay occurs upon the short incubation at 0°C (not shown). The experiments described above focused on the charge recombinations. In the following, we will address the forward electron transfer.

Figure 7 shows the flash-induced ΔI/I and its decay at 320 nm after the 2^nd^ flash given on dark-adapted WT*3-PSII (blue), WT*1-PSII (black) and ΔPsbJ-43H/PSII (red). The Q_A_^-^-*minus*-Q_A_ and Q_B_^-^-*minus*-Q_B_ difference spectra are similar and have their maximum amplitude at around 320 nm (Schatz and van Gorkom 1985). After the 1^st^ flash and the 3^rd^ flash, the S_1_ to S_2_ and S_3_ to S_0_ transitions also contribute significantly to the flash induced absorption changes at 320 nm (Lavergne 1991). In addition, the first flash is also complicated by the Q_A_^-^ to Fe^2+^ electron transfer (Boussac et al. 2011) with possible contribution in this spectral range of the Fe^3+^/Fe^2+^ couple (Sellés and Boussac, unpublished) and with the formation of the Tyr_Z_^•^ radical in inactive centers (with a flash spacing of 300 ms, the dead centers contribute mainly on the first flash due to the slow decay of Tyr_Z_^•^). For all these reasons, Figure 7 only shows the data after the second flash which is the easiest kinetics to analyze. In centers with Q_B_ oxidized in the dark-adapted sate the flash illumination forms the Q_A_^-^Q_B_ state and then the Q_A_Q_B_^-^ state. In these centers, the flash-induced ΔI/I does not decay and it is responsible for the remaining stable ΔI/I at the longest times. In dark-adapted centers with Q_B_^-^ present, the flash illumination forms the Q_A_^-^Q_B_^-^ state and then the Q_A_Q_B_H_2_ state. This reaction is at the origin of the decay in Figure 7 and this kinetics is similar in the 3 samples with a *t*_1/2_ of 2-3 ms. The similar apparent lag phase with a duration of ∼ 1 ms likely corresponds to the electron transfer between Q_A_^-^ and either Q_B_ or Q_B_^-^. This experiment shows that the forward electron transfer is not affected by the deletion of PsbJ.

**Figure 7:**
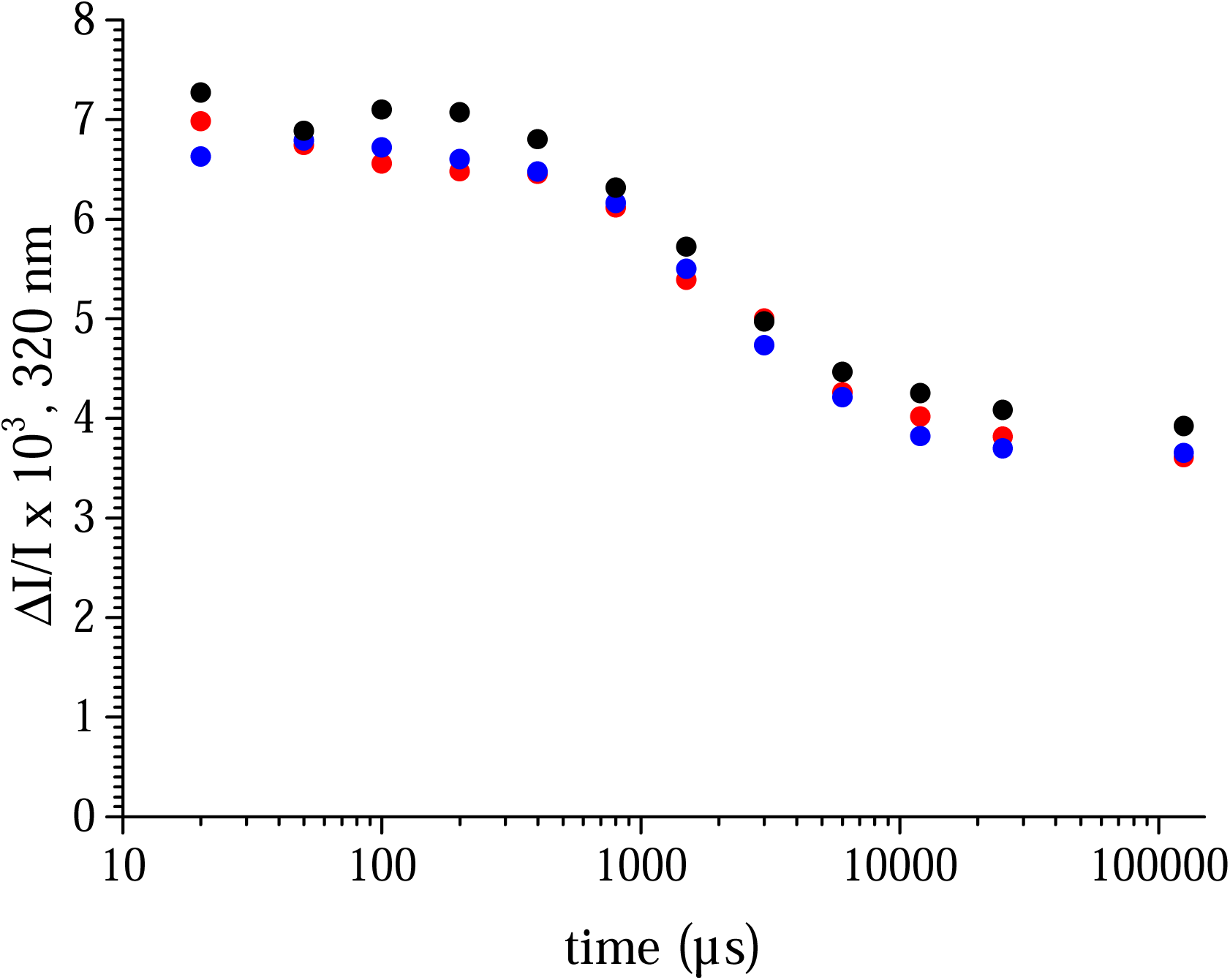
Time-courses of the ΔI/I changes at 320 nm after the 2^nd^ flash given on dark-adapted WT*1-PSII (black points), ΔPsbJ-43H/PSII (red points) and WT*3-PSII (blue data points). Flashes spaced 300 ms apart. Chl concentration adjusted to OD_673nm_=1.75.

Finally, Figure 8 shows EPR spectra in WT*1-PSII (black spectra) and ΔPsbJ-43H/PSII (red spectra) recorded with a magnetic field scale allowing a better observation of the quinone signals. Panel A shows the Fe^2+^Q_B_^-^ signal recorded in the dark-adapted samples. The signal is smaller in the ΔPsbJ-43H/PSII as mentioned previously and very slightly shifted. Panel B shows the spectra after a further illumination at 4.2 K. In centers with Q_B_^-^ present in the dark this 4.2 K illumination resulted in the formation of the Q_A_^-^Fe^2+^Q_B_^-^ signal that is larger in the WT*1-PSII. In the ΔPsbJ-43H/PSII, consequently to the larger proportion of centers with Q_B_ in the dark-adapted state, the 4.2 K illumination resulted in a larger proportion of Q_A_^-^Fe^2+^Q_B_ giving rise to the *g* = 1.9 signal between 3500 and 3700 gauss. Panel C shows the spectra recorded after an illumination at 4.2 K of samples in which a low amount of ferricyanide has been added (less than 100 µM) upon the dark adaptation to have the highest proportion of Q_B_ without the risk to oxidize the non-heme iron. Although there was still a low amount of Q_A_^-^Fe^2+^Q_B_^-^ signal in ΔPsbJ-43H/PSII, in both samples such a procedure resulted in the formation of a similar Q_A_^-^Fe^2+^Q_B_ characterized by the *g* = 1.9 signal. Finally, in Panel D the spectra were recorded after the addition of DCMU which resulted in the formation of Q_A_ ^-^Fe^2+^/DCMU in centers with Q_B_^-^ present in the dark. Two signals are observed in the two samples, the signal at *g* = 1.9 and a much broader ∼ 300 gauss-width signal reminiscent of a signal previously observed (Sedoud et al. 2011). The larger amplitude of these two signals in WT*1-PSII resulted of the higher concentration of Q_B_^-^ in the dark in this PSII.

**Figure 8:**
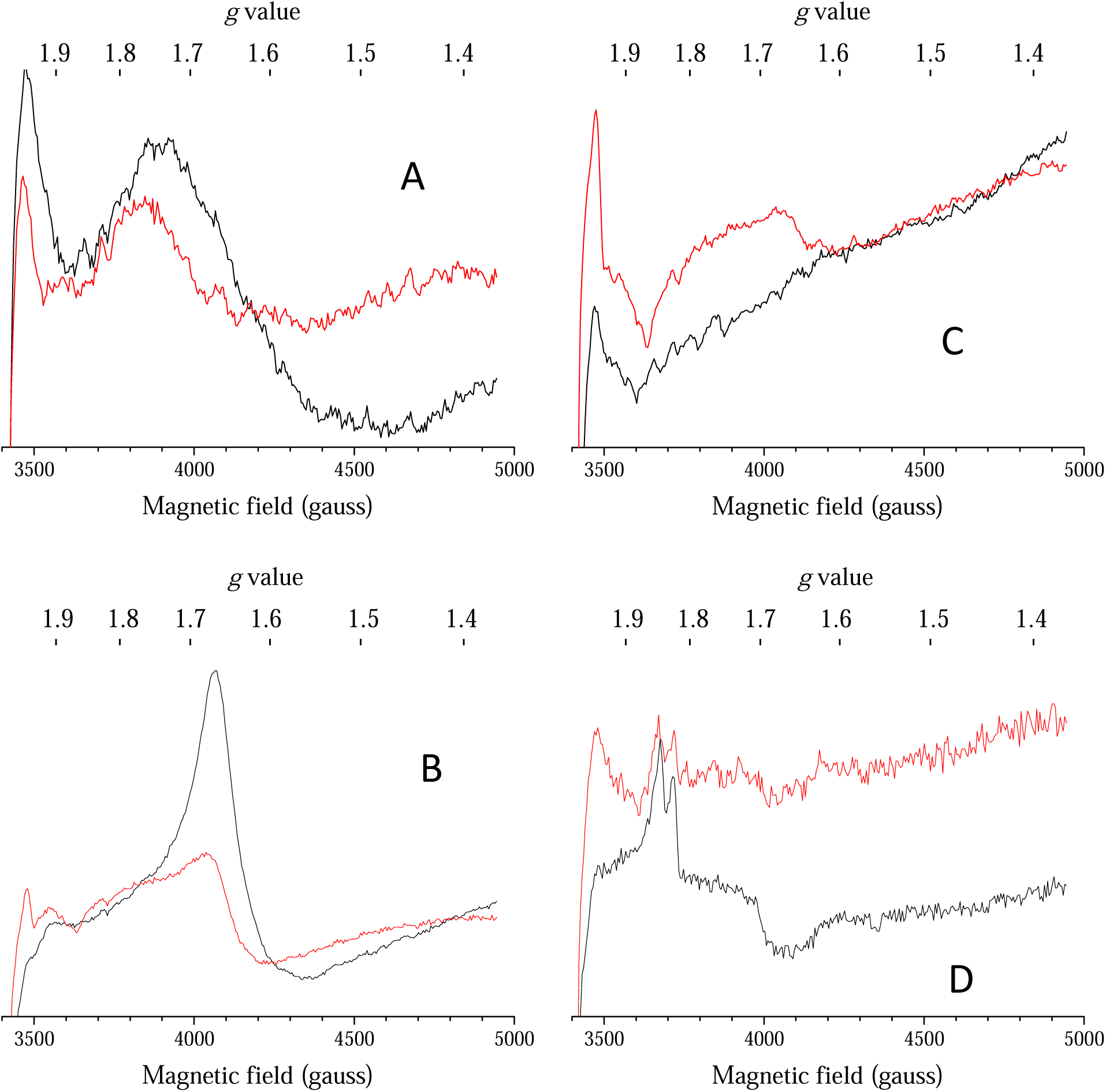
EPR spectra recorded in WT*1-PSII (black spectra) and ΔPsbJ-43H/PSII (red spectra). Panel A, the spectra were recorded after dark-adaptation for 1 hour at room temperature. Panel B, the spectra were recorded after a further illumination at 4.2 K. Panel C, the spectra were recorded after the addition of 100 µM of ferricyanide on dark-adapted samples followed by an illumination at 4.2 K. Panel D, the spectra were recorded after the addition of DCMU (100 µM) dissolved in ethanol to dark-adapted samples. The concentration was 1.1 mg Chl/mL. Instrument settings: temperature, 4.2 K except for Panel A in which T = 8.6 K; microwave frequency, 9.4 GHz; modulation frequency, 100 kHz; modulation amplitude, 25 G; microwave power, 20 mW.

## Discussion

In PSII lacking the PsbJ subunit it is possible to observe a proportion of the enzyme associated to polypeptides known to be assembly cofactors such as Psb27, Psb28 and Tsl0063, and which are not detected in the mature PSII, *e.g.* (Nowaczyk et al. 2006; Roose and Pakrasi 2008; Komenda et al. 2012; Liu et al. 2013; Huang et al. 2021; Zabret et al. 2021). The cryo-EM structure of an intermediate state was resolved (Zabret et al. 2021) with the interesting observation that the bicarbonate ligand of non-heme iron is replaced with a glutamate (glu241 of PsbD), a structural motif found in purple bacteria. Such a motif was further proposed to protect PSII from damage during biogenesis. Although the 3 proteins Psb27, Psb28 and Tsl0063 are possibility detected in the ΔPsbJ-43H/PSII studied here, the MALDI-TOF signals are so weak that we can reasonably consider that the purified PSII studied here is devoid of these assembly cofactors, see also (Sugiura et al. 2010b). The main peptides which are missing in this ΔPsbJ-43H/PSII are PsbY, PsbU and PsbV. Although PsbV is present in the ΔPsbJ-thylakoids (Sugiura et al. 2010b), the *g*_z_ value is lower than in intact PSII. This indicates (Roncel et al. 2003) that PsbV is not properly bound to PSII in thylakoids in the absence of PsbJ. Therefore, PsbJ very likely stabilizes the binding of the extrinsic peptides PsbU and PsbV and possibly of the trans-membrane α-helix PsbY as previously suggested (Zabret et al. 2021; Huang et al. 2021; Xiao et al. 2021). The ΔPsbJ-43H/PSII is essentially monomeric (Sugiura et al. 2010b) and in agreement with such a destabilization for PsbY, it was observed that although present, no electron density corresponding to PsbY was found in a crystal of monomeric PSII (Broser et al. 2010).

Despite the close proximity of PsbJ with the Q_B_ binding pocket, from the data in Figure 8, the lack of this subunit does not dramatically perturb, from an EPR point of view, any of the four states Q_A_Fe^2+^Q_B_^-^, Q_A_^-^Fe^2+^Q_B_, Q_A_^-^Fe^2+^Q_B_^-^ and Q_A_^-^Fe^2+^/DCMU. The quinone Q_B_ is present and DCMU binds in the Q_B_ pocket, with the same consequences, as in the wild type PSII. Nevertheless, the lack of strong structural modifications, the energetics is strongly modified. Since a great proportion of ΔPsbJ-43H/PSII has an intact Mn_4_CaO_5_ cluster, this allowed us to probe these changes by using thermoluminescence (Rutherford 1982; Johnson et al. 1995; Cser and Vass 2007; Rappaport and Lavergne 2009).

In WT*1-PSII, the TL peak corresponding to the S_2_Q_A_^-^/DCMU recombination is downshifted by ∼32°C, from 45°C to 13°C, when compared to the S_2_Q_B_^-^ recombination. According to a correspondence of 0.3-0.4°C/mV estimated by Rappaport and Lavergne (2009), see also (Cser and Vass 2007), this locates the energy level of the S_2_Q_A_^-^/DCMU state at least 80 mV (= 32/0.4) above the energy level of the S_2_Q_B_^-^ state in WT*1-PSII. If we assume that the *Em* of the Q_A_/Q_A_^-^ couple is increased by about 50 mV with DCMU bound as in plant PSII (Krieger et al. 1995), this would locate the energy level of the S_2_Q_A_^-^ state at least 80 + 50 = 130 mV above that of the S_2_Q_B_^-^ state in WT*1-PSII.

In ΔPsbJ-43H/PSII neither the S_2_Q_A_^-^/DCMU nor the S_2_Q_B_^-^ charge recombinations are detectable in the temperature range from −10°C to 70°C. The same reasoning done above for the WT*1-PSII would locate the energy level of the S_2_Q_A_^-^/DCMU state at least 200 mV (= 80/0.4) above the S_2_Q_B_^-^ state in ΔPsbJ-43H/PSII, 80°C being the lower limit for the difference in the TL peaks corresponding to the S_2_Q_A_^-^/DCMU and S_2_Q_B_^-^ charge recombinations in this sample.

We cannot totally discard the possibility that the release of PsbV affects the stability of S_2_ in the ΔPsbJ-43H/PSII. However, *Synechocystis* mutants lacking PsbV exhibit TL peaks at a slightly higher temperature than in the wild type (Shen et al. 1998) suggesting an increase of the S_2_ stability that is the opposite of what is seen here.

For analyzing the TL data, two extreme cases will be considered assuming that the effect of the deletion occurs mainly on the acceptor side. In the first one, the deletion of PsbJ would only affect the *Em* of the Q_A_/Q_A_^-^ couple whereas in the second case, only the *Em* of the Q_B_/Q_B_^-^ couple would be affected. We cannot discard a possible effect on both Q_A_ and Q_B_. In this case, the changes will apply on Q_A_ and Q_B_ but to a less extent on each of them.

In the first case, we will assume that in ΔPsbJ-43H/PSII the binding of DCMU also increases the *Em* of the Q_A_/Q_A_^-^ couple by ∼50 mV as supposed above for WT*1/PSII. Therefore, with an energy level of the S_2_Q_A_^-^/DCMU state at least 200 mV above that of the S_2_Q_B_^-^ state this would locate the energy level of the S_2_Q_A_^-^ state (without DCMU) at least 200+50=250 mV above the S_2_Q_B_^-^ state in ΔPsbJ-43H/PSII instead of 130 mV in WT*1-PSII. Although the value of 250 mV could be overestimated, this high value explains the lack of the S_2_Q_B_^-^ charge recombination experimentally observed in the TL experiment and the very fast S_2_Q_A_^-^/DCMU charge recombination. The decrease in the *Em* of the Q_A_/Q_A_^-^ couple could also explain the faster charge recombination observed in *Synechocystis* 6803 (Regel et al. 2001) in the absence of PsbJ if we are in conditions with a large proportion of the quinone pool fully reduced as often observed with whole cells.

In the second case, the *Em* of the Q_B_/Q_B_^-^ couple would reach a value disfavoring the electron coming back on Q_A_. However, the energy level of the S_2_Q_A_^-^/DCMU also need to be much higher in ΔPsbJ-43H/PSII than in WT*1-PSII to explain the lack of a TL signal in the presence of DCMU above −10°C. Since the peak in WT*1-PSII is observed at 13°C, a peak at a temperature below −10°C correspond to a decrease by at least ∼ −58 mV (–23/0.4) for the *Em* of the Q_A_/Q_A_^-^ couple in the presence of DCMU when compared to WT*1-PSII. Since PsbJ is close to the Q_B_ pocket and therefore close to the DCMU binding site, a DCMU effect on the *Em* of the Q_A_/Q_A_^-^ couple different in ΔPsbJ-43H/PSII than in plant PSII would not be unlikely. If we further push the reasoning assuming no change in the *Em* Q_A_/Q_A_^-^ couple in the ΔPsbJ-43H/PSII, the effect of the DCMU binding could be negligible. Indeed, the *Em* of the Q_A_/Q_A_^-^ couple with DCMU bound in the ΔPsbJ-43H/PSII would be the same than for the Q_A_/Q_A_^-^ couple in the absence of DCMU in WT*1/PSII. A higher *Em* of the Q_B_/Q_B_^-^ couple may also explain the lower O_2_ evolution under continuous illumination by the ΔPsbJ-43H/PSII (Sugiura et al. 2010b) by decreasing the efficiency of the electron transfer between Q_B_^-^ and the added quinone.

Alone, a low *Em* of the Q_A_/Q_A_^-^ couple in ΔPsbJ-43H/PSII is expected to increase the damages due to the repopulation of triplet states in the functional enzyme. For that reason we would favor the model in which only the *Em* of the Q_B_/Q_B_^-^ couple is affected (increased) in ΔPsbJ-43H/PSII. The increase of the Δ*Em* between Q_A_/Q_A_^-^ and Q_B_/Q_B_^-^, and without affecting the forward electron transfer, could contribute in a protection against the charge recombinations between the donor side and Q_B_^-^. Such a charge recombination was identified at the origin the damage by about 2 orders of magnitude higher than that induced by the same amount of energy delivered by continuous light (Keren et al. 1997) and a protection against it would favor the photoactivation process, see (Bao and Burnap 2016) for a recent review on photoactivation.

## Acknowledgments

This work has been in part supported by i) the French Infrastructure for Integrated Structural Biology (FRISBI) ANR-10-INBS-05, ii) the Labex Dynamo (ANR-11-LABX-0011-01), iii) EQUIPEX (CACSICE ANR-11-EQPX-0008), notably through funding of the Proteomic Platform of IBPC (PPI). MS was supported by the JSPS-KAKENHI grant in Scientific Research on Innovative Areas JP17H06435 and a JSPS-KAKENHI grant 21H02447.

## Abbreviations

Chl: chlorophyll
Chl_D1_/Chl_D2_: accessory Chl’s on the D1 or D2 side, respectively
DCMU: 3-(3,4-dichlorophenyl)-1,1-dimethylurea
PSII: Photosystem II
MES: 2–(*N*–morpholino) ethanesulfonic acid
P_680_: primary electron donor
P_D1_ and P_D2_: 
Chl monomer of P_680_ on the D1 or D2 side, respectively: Phe_D1_ and Phe_D2_, pheophytin on the D1 or D2 side, respectively
Q_A_: primary quinone acceptor
Q_B_: secondary quinone acceptor
Tyr_D_: redox active tyrosine 160 of D2
Tyr_Z_: redox active tyrosine 161 of D1
TL: thermoluminescence
WT*3: *T. elongatus* mutant strain containing only the *psbA*_*3*_ gene and a His_6_-tag on the C-terminus of CP43.
EPR: Electron Paramagnetic Resonance spectroscopy
MALDI-TOF: Matrix Assisted Laser Desorption Ionization - Time of Flight.

